# A workflow for rapid unbiased quantification of fibrillar feature alignment in biological images

**DOI:** 10.1101/2021.07.22.453401

**Authors:** Stefania Marcotti, Deandra Belo de Freitas, Lee D Troughton, Fiona N Kenny, Tanya Shaw, Brian M Stramer, Patrick W Oakes

## Abstract

Measuring the organisation of the cellular cytoskeleton and the surrounding extracellular matrix (ECM) is currently of wide interest as changes in both local and global alignment can highlight alterations in cellular functions and material properties of the extracellular environment. Different approaches have been developed to quantify these structures, typically based on fibre segmentation or on matrix representation and transformation of the image, each with its own advantages and disadvantages.

Here we present *AFT - Alignment by Fourier Transform*, a workflow to quantify the alignment of fibrillar features in microscopy images exploiting 2D Fast Fourier Transforms (FFT). Using pre-existing datasets of cell and ECM images, we demonstrate our approach and compare and contrast this workflow with two other well-known ImageJ algorithms to quantify image feature alignment. These comparisons reveal that *AFT* has a number of advantages due to its grid-based FFT approach. 1) Flexibility in defining the window and neighbourhood sizes allows for performing a parameter search to determine an optimal length scale to carry out alignment metrics. This approach can thus easily accommodate different image resolutions and biological systems. 2) The length scale of decay in alignment can be extracted by comparing neighbourhood sizes, revealing the overall distance that features remain anisotropic. 3) The approach is ambivalent to the signal source, thus making it applicable for a wide range of imaging modalities and is dependent on fewer input parameters than segmentation methods. 4) Finally, compared to segmentation methods, this algorithm is computationally inexpensive, as high-resolution images can be evaluated in less than a second on a standard desktop computer. This makes it feasible to screen numerous experimental perturbations or examine large images over long length scales.

Implementation is made available in both MATLAB and Python for wider accessibility, with example datasets for single images and batch processing. Additionally, we include an approach to automatically search parameters for optimum window and neighbourhood sizes, as well as to measure the decay in alignment over progressively increasing length scales.

## 1 Introduction

Measuring the anisotropy of features in biological images is of increasing interest as the degree of alignment can inform both the underlying cellular behaviours and material properties of the sample. For example, different cell types have unique emergent capacities to align in culture, controlled by cellular packing and geometry (Duclos et al., 2014, 2017). When cells are exposed to external forces such as cyclical stretch, they tend to align their cytoskeleton perpendicular to the axis of stretch (Standley et al., 2002; Yoshigi et al., 2005; Livne et al., 2014). Subcellular cytoskeletal networks can also spontaneously organise in response to the stresses of their environment (Gupta et al., 2015, 2019) or changes in biochemical signalling (Ridley and Hall, 1992). Similar changes in cytoskeletal architecture can be seen in reconstituted protein systems (Falzone et al., 2013; Linsmeier et al., 2016). Additionally, the extracellular matrix (ECM) also has an inherent capacity to align in different tissues or pathologies, such as cancer (Ouellette et al., 2021) or tissue fibrosis (Park et al., 2020; Mascharak et al., 2021), which is thought to alter the ECM network mechanical properties. These examples illustrate the variety of environments where alignment of features reveals important biological properties. As such, it is necessary to develop approaches to efficiently quantify anisotropy of features across a range of length scales, from subcellular organisation to tissue level alignment.

A number of different approaches have been developed to quantify the alignment of image features. These methods can be categorised based on the type of algorithm used to highlight features and subsequently quantify anisotropy. Fibre segmentation tools, such as the Fiji Ridge Detection plugin (Lindeberg, 1998), the curvelet-based CurveAlign/CT-FIRE suite (Bredfeldt et al., 2014a,b; Liu et al., 2017, 2020), and Filament Sensor (Eltzner et al., 2015), provide individual fibre information. Tools based on representation/transformation of the image, such as the Fiji plugins OrientationJ (Rezakhaniha et al., 2012) and FibrilTool (Boudaoud et al., 2014) or the CytoSpectre suite (Kartasalo et al., 2015), supply overall fibre alignment information. Hybrid tools, such as TWOMBLI, which exploit a combination of approaches (i.e., fibre segmentation followed by matrix representation of the image) have also been recently made available (Wershof et al., 2021).

Many image transformation algorithms rely on Fourier transformation of the image and exploit its frequency space representation to obtain alignment information (Chaudhuri et al., 1987; Pourdeyhimi et al., 1997; Marquez, 2006; Sander and Barocas, 2009; Goldyn et al., 2010; Kartasalo et al., 2015; Clemons et al., 2018). These algorithms tend to be computationally efficient (Sander and Barocas, 2009) and insensitive to the image background noise (Liu et al., 2020), which is desirable for rapidly processing large volumes of experimental data (Püspöki et al., 2016). Another desirable trait for an alignment detection tool is its flexibility in terms of size and location of alignment patterns (Püspöki et al., 2016), which is handled well by the only open-source software available in this category (CytoSpectre (Kartasalo et al., 2015)) due to a power spectrum analysis of the image’s discrete Fourier transform.

One feature missing from the available tools in the literature is an analysis of the alignment length scale. Depending on the scientific question of interest, one may want to measure alignment over a small region of the image (e.g., local cytoskeletal features inside cells) or over a broad length scale (e.g., ECM organisation in whole tissues). Additionally, when comparing experimental datasets, it is important to understand the precise length scale over which the observed alignment is significant and determine the spatial decay of the anisotropy, which will allow for better interpretation and analysis of results (e.g., is the alignment of a region of interest a subcellular or supra-cellular phenomenon). Here we explain the implementation of an open-source alignment quantification algorithm, *AFT - Alignment by Fourier Transform*, that can be run on a variety of biological images and has a number of advantages over pre-existing approaches. This FFT-based quantification is rapid and computationally efficient, flexible to different types of microscopy images and samples, easy to use with a low number of parameters, and importantly allows for a user-defined length scale over which to measure feature anisotropy.

## 2 Materials and Methods

Sample datasets of microscopy images were used to illustrate the analysis. In particular, cultured fibroblasts transfected with GFP-actin, fibroblasts fixed and stained for filamentous actin (F-actin, phalloidin), fixed cell-derived matrices immunostained for actin (before decellularisation) and fibronectin (after decellularisation), and second harmonic generation imaging of tissue samples were utilised. Fixed actin images were acquired at different magnifications using either single cells or cells at confluence, to allow for evaluation of the length scale of alignment.

Fixed samples were imaged using a Zeiss LSM 880 equipped with a 40x NA 1.3 Plan-Apochromat oil objective or 63x NA 1.4 Plan-Apochromat oil objective. Decellularised matrices were imaged using a Zeiss LSM 880 equipped with 63x NA 1.4 Plan-Apochromat oil. For second harmonic generation imaging, tissue sections were imaged using a Zeiss LSM 7MP equipped with a 20x NA 1.0 water immersion objective. Live cell imaging was performed on a 3i Marianas Imaging System consisting of a Zeiss Axio Observer 7 inverted microscope attached to a Yokogawa W1 Confocal Spinning Disk using a 63x NA 1.4 Plan-Apochromat oil objective.

### 2.1 Pre-processing

Image pre-processing is not required for this analysis, but can be used to additionally highlight the fibrillar features. In the context of this work, pre-processing was performed only for the cell-derived matrix images, as their surface is often not flat resulting in uneven signal across the field of view. To account for this, contrast was enhanced in Fiji (0.35% saturated pixels) and subsequently a local contrast adjustment was performed (CLAHE plugin, default parameters). All images shown are maximum projections of Z-stack acquisitions, except for the live cell images which represent a single confocal slice.

### 2.2 Comparison with other available alignment tools

The performance of the algorithm presented in this work was compared with OrientationJ (Rezakhaniha et al., 2012) and TWOMBLI (Wershof et al., 2021), by using two 10-image samples of fibronectin-stained cell-derived matrices that were either isotropic or anisotropic. To access the alignment vector field for OrientationJ, a Fiji macro was designed to batch run the OrientationJ Vector Field plugin. This operation was either performed on its own or preceded by fibre segmentation by the Ridge Detection plugin (Lindeberg, 1998), this second approach mimicking the TWOMBLI workflow (hybrid method). To obtain a comparable spatial resolution of the vector field, a window of 100 *px* with 50% overlap was used to run the FFT analysis, and a matching local window *σ* of 100 *px* with a grid size of 50 was employed in OrientationJ. The Ridge Detection parameters were set to: *line width =* 20 *px*, *high contrast =* 120, *low contrast =* 0, *sigma =* 6.27, *lower threshold =* 0, *upper threshold =* 0.17, *minimum line length =* 10 *px*. These parameters were selected to match the ones assigned by the TWOMBLI macro when calling the Ridge Detection plugin (see below). From the acquired vector fields, an order parameter over neighbourhoods of 5*x* vectors was calculated and compared for the three approaches.

A second comparison was based on coherency, the metric traditionally used for global alignment output by both OrientationJ and TWOMBLI. Coherency of the FFT was measured on windows of 100 *px* with 50% overlap and averaged to obtain a median value representing the global alignment score for each image. To obtain the coherency score from OrientationJ, a macro was designed to batch process the images using the OrientationJ Dominant Direction plugin. TWOMBLI was run on the sample data as per the developer instructions. A parameter optimisation step was performed on test images and the obtained parameter file was used for batch processing (*line width =* 20 *px*, *curvature window =* 150 *px*, *branch length =* 10).

Computational time was measured on a 3.6 GHz Quad-Core Intel Core i7 machine with 32 GB memory, with suitable functions in MATLAB and ImageJ macro language. Only the effective analysis time for a 10-image sample (1 MB per image) was taken into account, excluding the input and parameter upload. Timings for OrientationJ refer to the designed macro, no manual handling of data was involved.

### 2.3 Statistical analysis

Statistical analysis was performed in Prism (Graphpad, v9). Mann-Whitney two-tailed tests were used to compare metrics on isotropic vs. anisotropic samples. Significance is reported in figure panels as follows: ‘****’ for p-values lower than 0.0001, ‘***’ lower than 0.001, ‘**’ lower than 0.01, ‘*’ lower than 0.05, ‘ns’ otherwise.

### 2.4 Implementations

This approach builds upon previous work employed in (Aratyn-Schaus et al., 2011; Cetera et al., 2014; Fernandes et al., 2020) and adds features such as periodic decomposition to improve the accuracy of the FFT angle determination along with the ability to search for the optimal length scale for comparison. The current implementation is available at https://github.com/OakesLab/AFT-Alignment_by_Fourier_Transform, both in MATLAB (Mathworks, v2018b) and Python languages. The MATLAB version is provided with a simple user interface for inputting parameters, while the Python version is presented as a set of Jupyter notebooks. The code is divided into two separate suites. The first suite can be used to perform the alignment quantification on a sample of images containing fibrillar structures. The second suite can be used to run a search to optimise the analysis parameters, with the aim of maximising differences between two samples, i.e., locating the length scale for which the difference in alignment between two samples is greatest. Further code documentation is provided in the repository.

## 3 Results

### 3.1 Measuring alignment

The algorithm is inspired by the general approach of Particle Image Velocimetry (Willert and Gharib, 1991), where the image is broken down into a series of windows that are analysed independently to create a vector field that represents the entire image. Windows are analysed in frequency space to reveal information about both the fibrillar structure and local alignment as illustrated in Fig 1 using filamentous actin images. If the image contains aligned features in the real space, the corresponding FFT in the frequency domain will be asymmetrically skewed, with the direction of skew orthogonal to the original feature orientation (Fig 1AB). This process can then be repeated on each successive window across the image, resulting in a vector field that represents the local alignment in the image. A detailed protocol for determining the local alignment of each window and the order parameter of global alignment follows.

**Figure 1:**
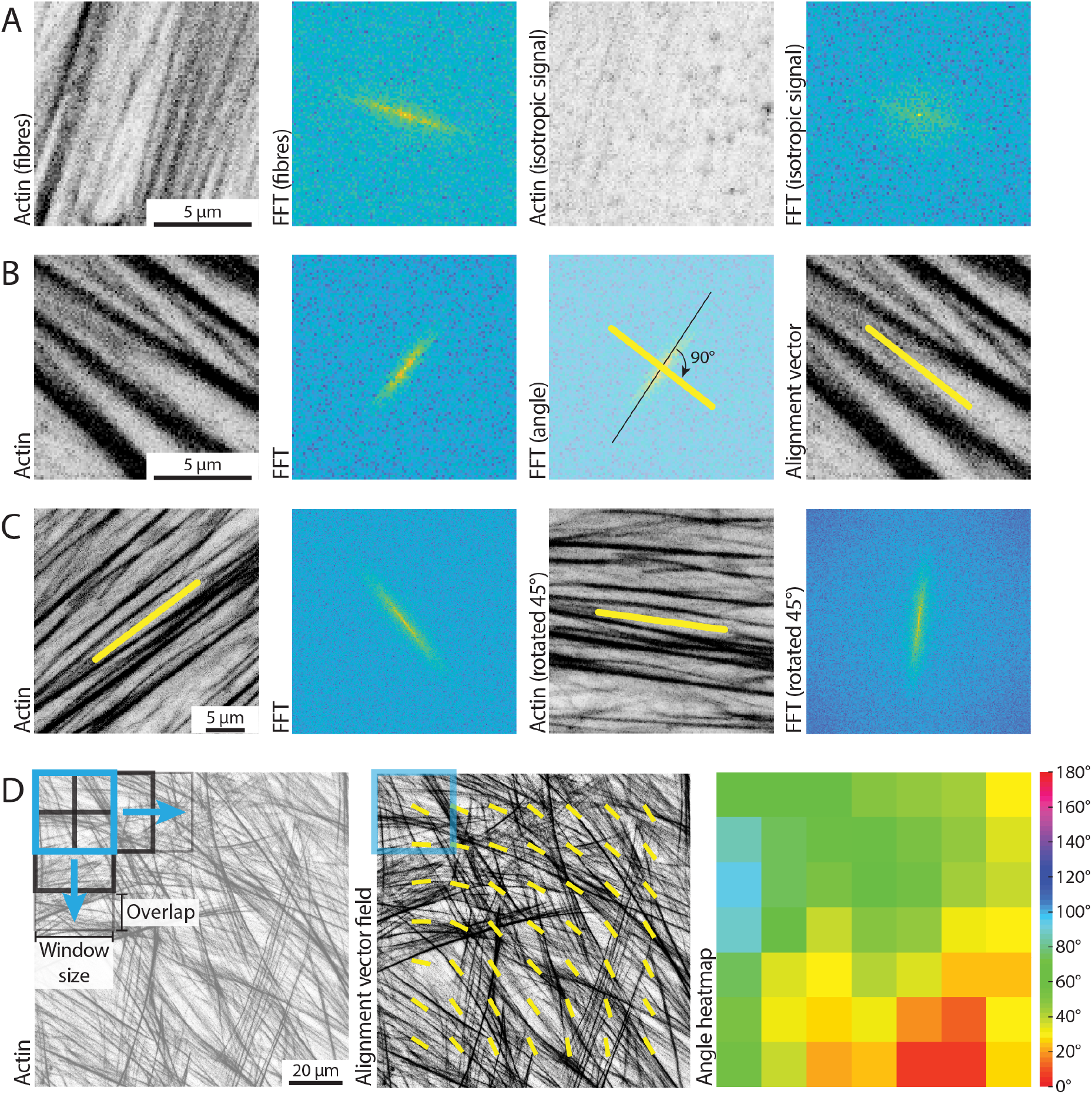
Using FFTs to measure local alignment of biological images. **(A)** Small windows of a microscopic image are considered, containing either aligned filamentous actin fibres, or isotropic signal. The 2D Fast Fourier Transform (FFT) shows an elongated (i.e., skewed) shape if the image in the real space contains aligned fibrillar features, and displays a more round shape otherwise. Scale bar, 5 *μm*. **(B)** A small window of a microscopic image containing aligned filamentous actin fibres is considered. Its FFT shows a predominant skewness (black line) of an angle that is orthogonal to the direction of the fibres in the real space image. The obtained alignment vector (yellow line) can be overlaid to the original image to highlight the measured fibre orientation. Scale bar, 5 *μm*. **(C)** The alignment vector is calculated for an image containing aligned actin fibres by exploiting the characteristics of the FFT. The same image, rotated by 45°, is analysed, to demonstrate robustness of the method to rotation. Scale bar, 5 *μm*. **(D)** A square representing the window size for analysis (250 *px*, light blue) is overlaid on a filamentous actin image. The black squares represent subsequent instances of such a window with the defined overlap, as the image is scanned during the analysis to calculate local alignment vectors (yellow). The heatmap shows a different representation of the alignment vector field, with a wrapped colour scale ranging from 0° to 180°. Scale bar, 20 *μm*.

#### 3.1.1 Local alignment

Square windows of *n x n* pixels are chosen from the original real space image and a 2D FFT is performed. Typically, *n* is chosen as odd to ensure that the zero-order frequency component, which is always the greatest in magnitude, is situated at the centre of the FFT. FFTs of non-periodic signals are often plagued by strong horizontal and vertical components, appearing as a cross, due to the mismatch of intensity in the images at the edges. To avoid this effect, we break down the image into its smooth and periodic components following the approach of Moisan (2011). We use the periodic component of this decomposition to take the FFT and then take its norm to only deal with real numbers. The resulting image is then multiplied by a mask of diameter *n/*2 to capture all the relevant high frequency components and to ensure the sample is symmetric. The central image moments of the masked FFT are then calculated as

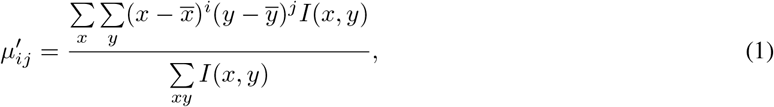

where *x* and *y* represent the position and *I* is the magnitude of the FFT norm. We next construct the covariance matrix of the image as

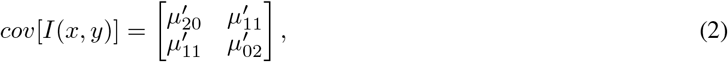

which has eigenvalues of

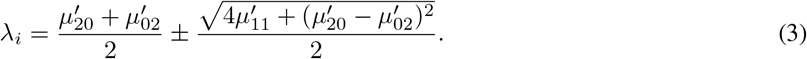

The orientation of the FFT can thus be calculated by

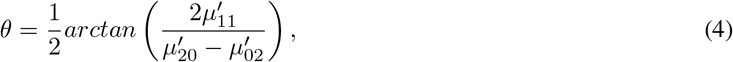

and the eccentricity, a measure of how oblong the FFT is, as

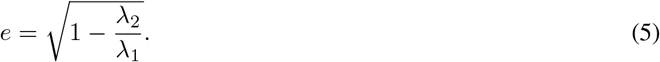

The orientation of the features in the real image is orthogonal to the orientation of the FFT, and thus we apply a 90° rotation (Fig 1B). Orientation vectors do not have a polarity, and thus angles of 0° or 180° are considered equivalent. Therefore, all angles are mapped to a range of 0° to 180° (Fig 1D). This methodology is robust to rotational transformations and does not suffer from the inherited bias of FFT for vertical and horizontal components (Fig 1C).

#### 3.1.2 Order Parameter Calculation

Both the size of the window and the degree of overlap between the windows is customisable (Fig 1D). To measure how aligned fibrillar features are within adjacent windows, we define an order parameter by correlating the directionality of vectors within the neighbourhood of customisable size. Specifically, the order parameter can be calculated as

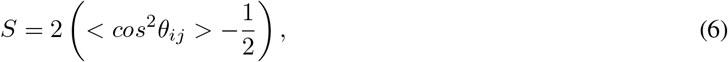

with *θ_ij_* representing the angle between the orientation of a central reference vector and its neighbours (Fig 2A). The order parameter can take values between −1 and 1 (Fig 2A), with 1 representing perfect alignment (i.e., all vectors in the neighbourhood have the same orientation as the reference vector), 0 representing random orientation, and −1 representing opposite alignment (i.e., all vectors in the neighbourhood are pointing in the orthogonal direction compared to the reference vector). While a number of different formulations of the order parameter have been used previously, in the context of the current analysis, the chosen order parameter normally ranges between 0 (random) and 1 (perfectly aligned), as fibre polarity is not taken into account and the neighbourhood area is kept relatively local (Fig 2B).

To calculate an overall alignment score for each image, we define a neighbourhood size. The previously calculated vector field is then split into all possible overlapping neighbourhoods of such size. The order parameter is subsequently calculated for each neighbourhood, and the overall order parameter for the image is defined as the median value (Fig 2B).

**Figure 2:**
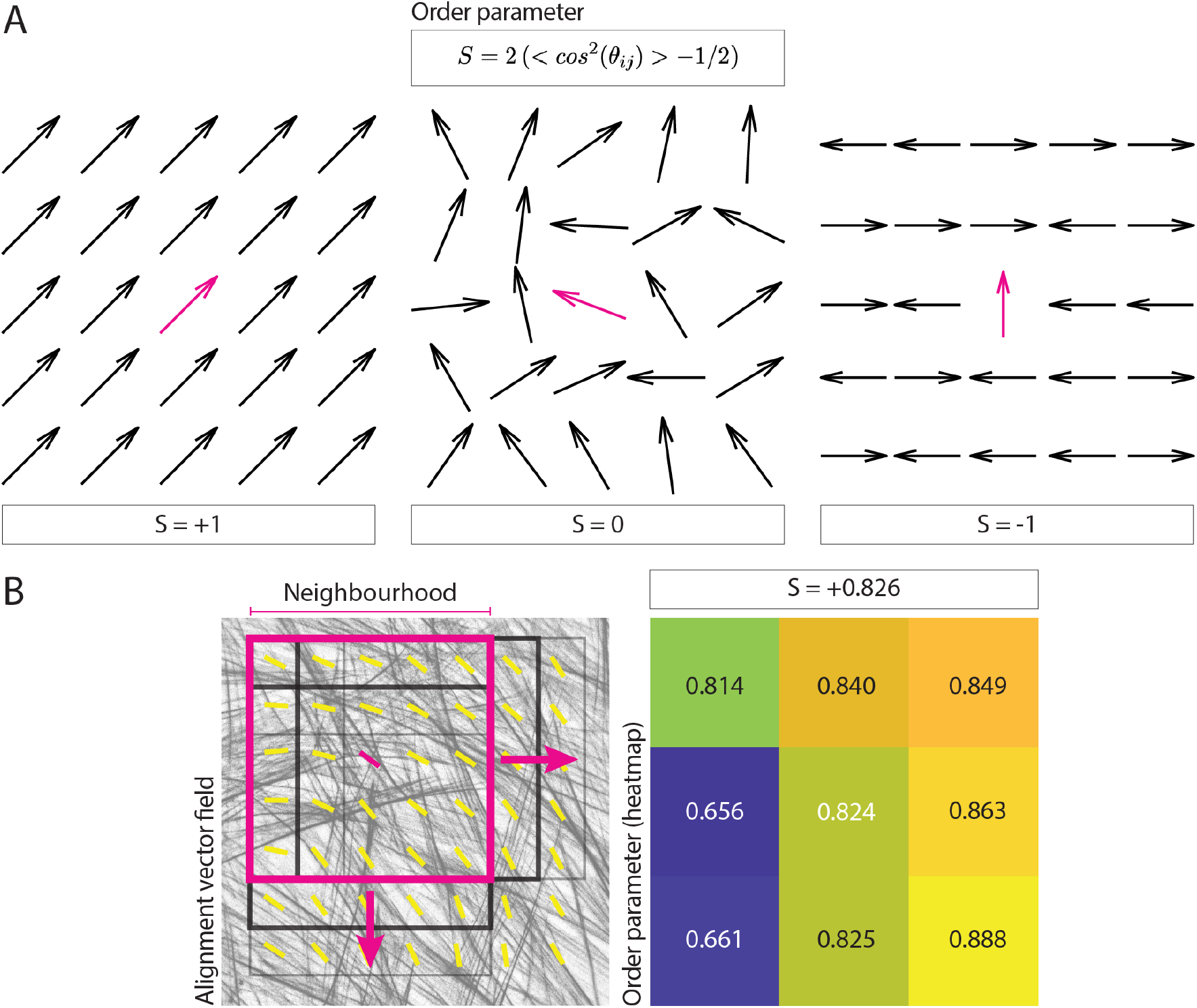
Calculating an image order parameter. **(A)** The equation for the order parameter used to assign an alignment score for each neighbourhood is shown, together with examples. The order parameter takes a value of 1 for complete alignment, 0 for random alignment, and −1 for orthogonal alignment. **(B)** A square representing a neighbourhood of 5*x* vectors (magenta, neighbourhood radius of 2*x* vectors around the central reference) is overlaid on an actin image and its corresponding alignment vector field. The black squares represent subsequent instances of such neighbourhood, as the vector field is scanned during the analysis to obtain local alignment scores. The resulting order parameters for each neighbourhood are reported in the heatmap, together with their median value, representing the output of the analysis.

### 3.2 Investigating the optimal alignment length scale between two samples

When comparing two samples displaying different degrees of alignment, it may be of interest to evaluate the length scale for which this difference is greatest. The approach described above allows for this, as it is possible to evaluate the order parameter using a range of window and neighbourhood sizes which correspond to different length scales (i.e., each permutation of window and neighbourhood size define a specific length scale). By comparing the obtained alignment scores between the two experimental conditions for the different parameter permutations, it is possible to investigate the precise length scale at which the difference between the two samples is most pronounced. The comparison can be carried out by looking at the order parameter difference between the two samples or by running statistical tests and analysing the resulting p-values.

We tested this parameter search approach on actin images of cultured fibroblasts with different degree of isotropy (Fig 3A, 10 images for each sample), for window sizes ranging from 25 to 325 *px* and neighbourhood radii ranging from 1*x* to 38*x* vectors (Fig 3BC). We performed the comparison in alignment scores for each window/neighbourhood size permutation by quantifying the difference between the sample order parameters (Fig 3B) or by calculating the p-value of a Mann-Whitney statistical test between the two populations (Fig 3C).

**Figure 3:**
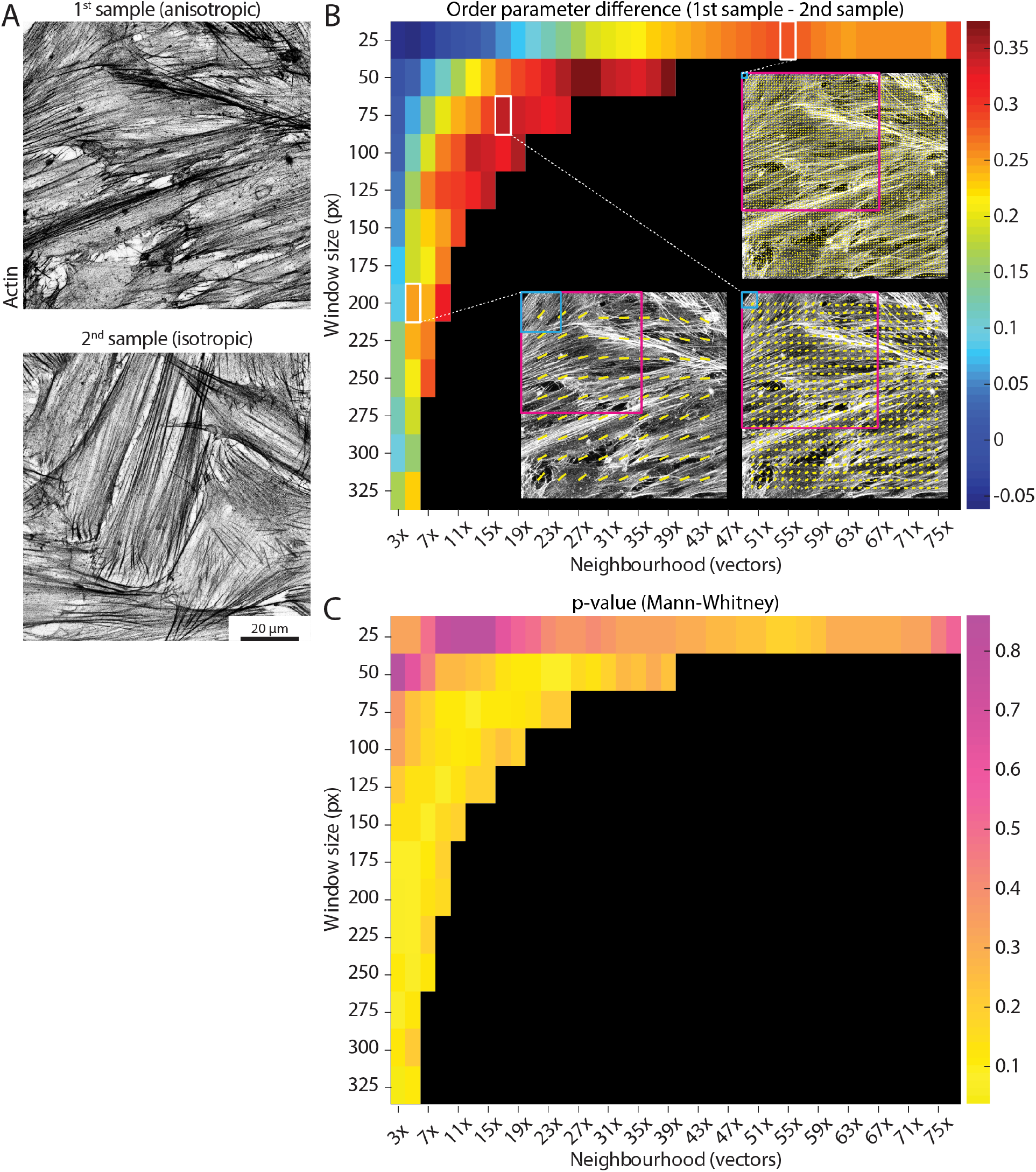
Determining the optimal length scale difference. Example actin images taken from two separate samples and displaying different degrees of alignment (higher on the left, more anisotropic; lower on the right, more isotropic). Scale bar, 20 *μm*. **(B)** The parameter search runs the analysis for a range of window sizes (y-axis, from 25 *px* to 325 *px*) and a range of neighbourhood sizes (x-axis, from 3*x* to 77*x* vectors, where the neighbourhood size is defined as 2 *neighbourhood radius +* 1). Each permutation of window and neighbourhood represent a length scale. Each coloured square shows the difference in the median order parameter between the anisotropic sample and the isotropic sample (mean values over 10 images for each sample) for a specific pair of window and neighbourhood sizes. Small window sizes paired with small neighbourhoods (upper left corner) display noisy output, due to the limited amount of image information available to process. Three example vector fields are shown for a window size of 25 *px* and neighbourhood of 55*x* vectors (corresponding to 92 *μm*), 75 *px* and 17*x* vectors (89 *μm*), and 200 *px* and 5*x* vectors (79 *μm*), with the light blue and magenta squares showing the size of the window and neighbourhood respectively. Despite relatively similar neighbourhood sizes, the output varies depending on the size of the examined window. Reading the graph horizontally from left to right for each window size up to 150 *px*, it is possible to note how an optimal length scale of 90 *μm* is detected (i.e., a peak difference value is displayed), likely correlated to relevant biological dimensions (e.g., cell size). **(C)** A similar graph to the one in (B) is shown for the p-value of Mann-Whitney tests between the two samples for all the combinations of window and neighbourhood sizes. In this case, greater differences are represented by lower values (yellow).

It should be noted that small windows paired with small neighbourhoods display noisy output (Fig 3BC), as the corresponding FFT for such small windows will tend towards low eccentricity values (i.e., increasing the likelihood of spurious vectors, Fig 1A). Interestingly, when comparing two samples with known differences in alignment (Fig 3A), it can be observed how the difference in order parameter between the two samples displays a peak when keeping the window size constant and increasing the neighbourhood (Fig 3B). This suggests a well-defined optimum length scale where a clear difference between the two samples occurs, and it is likely to be correlated with relevant dimensions in the sample (e.g., cell size). It is important to note that the optimal length scale will likely exist for a range of different window/neighbourhood pairs, which together will define a similar local area. In the example shown in Fig 3B, the order parameter difference peaks for a length scale of ~ 90 *μm*, as calculated by multiplying the window size by the neighbourhood size for pairs displaying high difference values. This suggests that this length scale should be investigated further to evaluate the biological relevance of features of such size.

### 3.3 Evaluating the length scale decay in alignment

A similar approach to the parameter search can be employed to look at how the alignment decays over increasingly larger length scales, by progressively enlarging the size of the region over which the alignment is measured. To this aim, the window size over which the angle vector field is calculated is kept constant, while progressively increasing the neighbourhood over which the order parameter is evaluated.

We exemplified the alignment decay analysis by using 1 ~ *mm*^2^ tilescan images of cell-derived matrices (i.e., matrices produced *in situ* by the cells) stained for F-actin and fibronectin (before and after decellularisation, respectively, Fig 4A). We first calculated the angle vector field for both actin and fibronectin using the FFT approach for a window size of 250 *px* and overlap of 50% (Fig 4B). Subsequently, we calculated the median order parameter over neighbourhoods of increasing radii, ranging from 1*x* to 21*x* vectors, corresponding to 50 *x* 50 *μm* to 1050 *x* 1050 *μm* in the original image (Fig 4ACD). By plotting the median order parameter over the neighbourhood size, it is possible to observe the alignment decaying from the local to the global length scale (Fig 4D). As expected, actin and fibronectin displayed a similar decay in alignment as cells are responsible for depositing and remodelling the ECM in this experiment.

**Figure 4:**
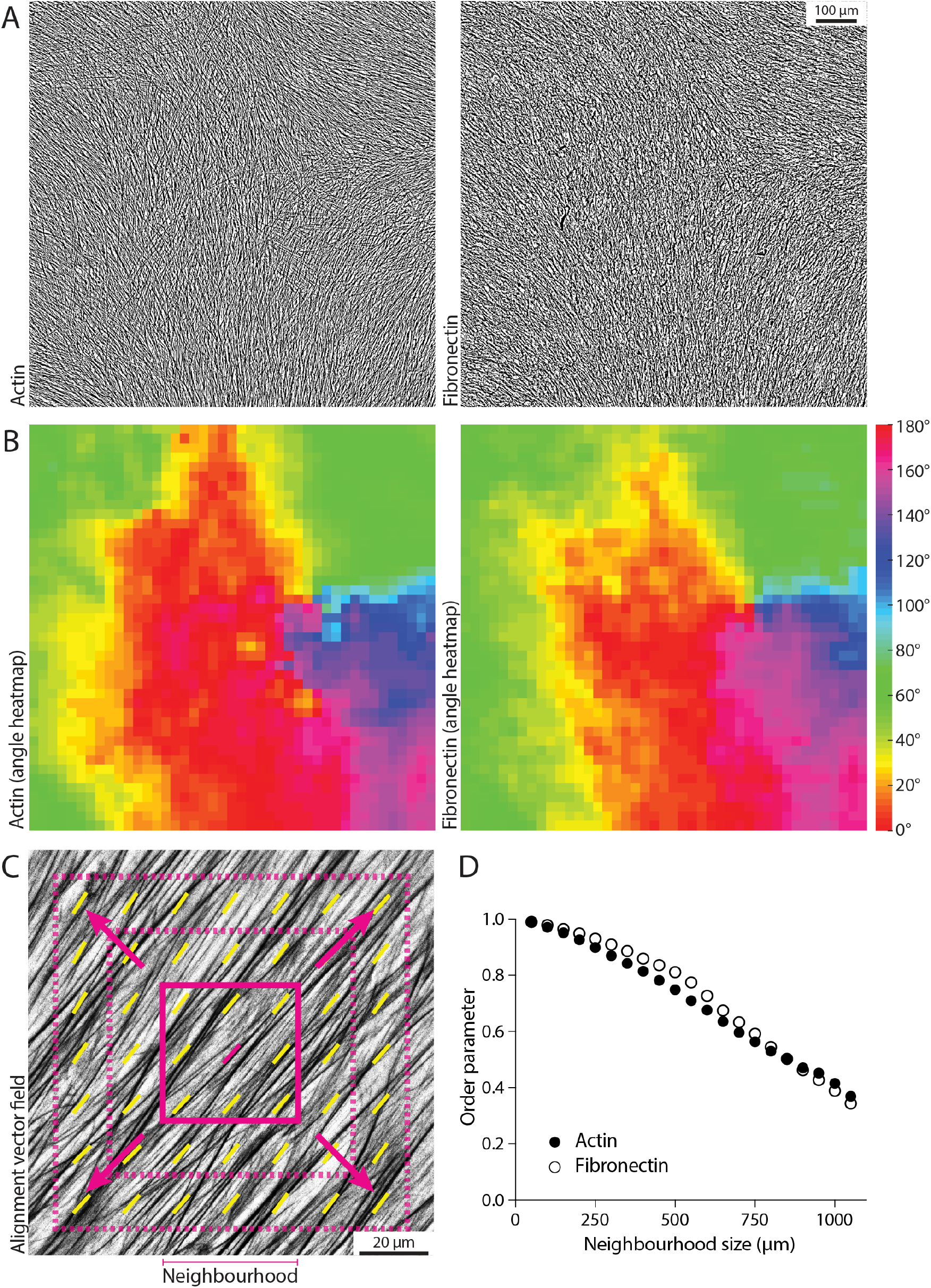
Measuring the decay in order parameter as a function of length scale. **(A)** Example tilescan images about 1 *mm*^2^ in size for actin and fibronectin on the same sample (fibroblast-derived matrix). Scale bar, 100 *μm*. **(B)** Angle heatmap showing the alignment vector field for the images in (A), using a window size of 250 *px* and 50% overlap. Actin and fibronectin display similar alignment. **(C)** Schematic of the alignment decay analysis, where the window size is maintained constant while increasing the size of the neighbourhood (i.e., including more and more vectors in the order parameter calculation to evaluate alignment from the local to the global scale). The solid magenta square represents the starting neighbourhood, which then increases in size at each iteration (dashed magenta squares). **(D)** The calculated order parameter for actin and fibronectin for the images in (A) is graphed for a window size of 250 *px*, for each neighbourhood ranging from 50 *μm* (3*x* vectors, neighbourhood radius of 1*x* vector) to 1050 *μm* (43*x* vectors, neighbourhood radius of 21*x* vectors). Alignment decays in a similar manner when going from local (small neighbourhood size) to global (large neighbourhood size) length scales.

### 3.4 Filtering input for alignment analysis

Some biological images might display regions that are either blank (i.e., little to no signal, lack of image features), or isotropic (i.e., no obvious fibrillar features). In certain applications, such regions should be excluded from the alignment analysis, as images containing noisy or blank areas will affect the output order parameter. By exploiting our window-based algorithm, it is possible to implement optional filtering of features to automatically exclude regions with such characteristics from the analysis.

To filter out blank regions, a threshold can be set on the mean intensity for each window, as demonstrated on second harmonic generation imaging of tissue samples (Fig 5A). In this example, windows that are considered background have a mean intensity signal lower than 20 (with intensity values ranging from 0-black to 255-white). This value can be set as a threshold: windows displaying mean intensity values lower than the threshold will be ignored during the calculation of the angle vector field (and subsequent alignment score, Fig 5B).

**Figure 5:**
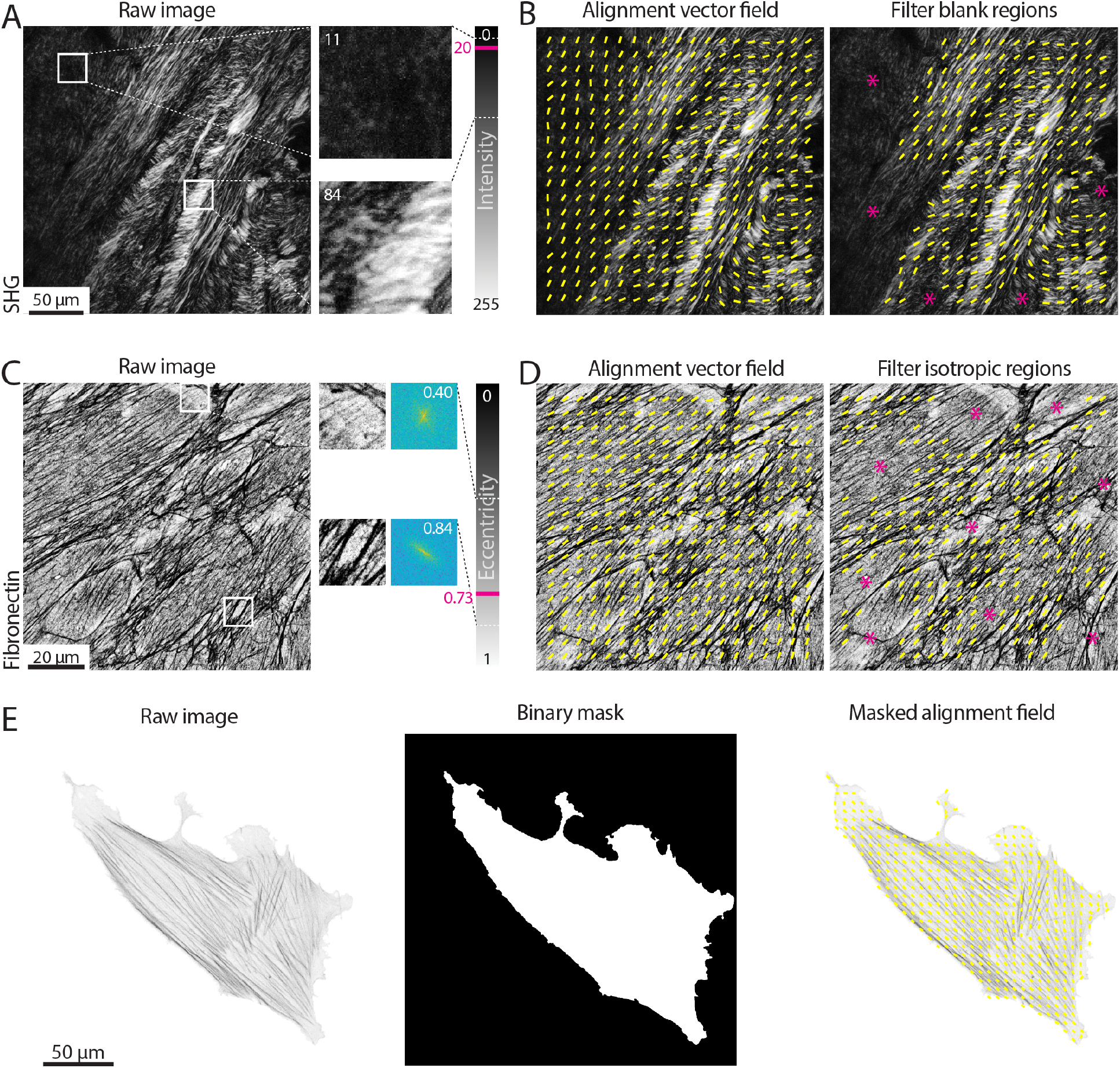
Filtering vector fields based on intensity and eccentricity. **(A)** A tissue section second harmonic generation (SHG) image is shown. Two regions with the same size of the window used for the alignment analysis (100 *px*) are enlarged and their mean intensity value measured. The intensity of an 8-bit image can vary from 0 (black) to 255 (white). The area on the top left contains little to no signal and it is considered background (mean intensity of 11); the area on the right contains many bright pixels, for a mean intensity of 84. A threshold value of 20 is set, meaning that all windows with mean intensity lower than such values will be considered as blank (i.e., background) and excluded from the analysis. Scale bar, 50 *μm*. **(B)** The alignment vector field (yellow) is overlaid on the raw SHG image, either without filtering for blank regions (left) or filtering with a threshold on the mean intensity of 20 (right). Magenta asterisks highlight regions excluded from the analysis. **(C)** An immunostaining for fibronectin in cell-derived matrices is shown. Two regions with the same size of the window used for the alignment analysis (100 *px*) are enlarged and their 2D Fast Fourier Transform (FFT) displayed. The FFT eccentricity can range from 0 (circular) to 1 (highly elongated). The region on the top lacks clear aligned fibres, resulting in a more uniform FFT (eccentricity of 0.40); the area on the bottom has aligned fibrillar features, leading to a highly skewed FFT (eccentricity of 0.84). A threshold value of 0.73 is set, meaning that all windows with eccentricity lower than such values will be considered as isotropic and excluded from the analysis. Scale bar, 20 *μm*. **(D)** The alignment vector field (yellow) is overlaid on the raw fibronectin image, either without filtering for isotropic regions (left) or filtering with a threshold on the eccentricity of 0.73 (right). Magenta asterisks highlight the regions excluded from the analysis. **(E)** The actin in a single cell is shown. If one wants to exclude the background from the analysis, a binary mask can be provided highlighting the regions of the image to be analysed (window size 50 *px* with 50% overlap). Scale bar, 50 *μm*.

Regions in which image features display isotropic organisation can also be excluded by setting a threshold on the eccentricity of the FFT calculated for a given window, as shown for cell derived matrices immunostained for fibronectin in Fig 5C. The FFT eccentricity can be used as a measure of its skewness (Fig 1A): regions containing oriented fibres will display a more elongated FFT, hence higher eccentricity (values closer to 1); regions where no fibres can be detected, or where there are multiple fibres without a clear orientation, will display a more homogeneously shaped (i.e., round) FFT, hence lower eccentricity (values closer to 0). In the example shown in Fig 5C, a threshold on the eccentricity was set to 0.73, allowing for the exclusion of isotropic regions from the angle vector field calculation (Fig 5D).

Finally, it is possible that only a specific region within an image is to be analysed, as in the case of actin in single cells (Fig 5E). A binary image can be used to mask the angle vector field, in order to only consider a given area of the original input.

### 3.5 Extracting kinetics of organisation from live imaging

Many biological processes involve the evolution of architectures and organisations over time. By measuring an order parameter for each frame of a time series it is possible to extract the kinetics of alignment in response to perturbations. As an example, we transfected an NIH 3T3 fibroblast with GFP-actin and imaged the cell before, during, and after treatment with the Rho kinase inhibitor Y-27632 (Fig 6). Inhibition of Rho kinase leads to a reduction in myosin activity and thus a dissolution of stress fibres (e.g., a loss of fibrillar structures; Fig 6A). As a test case, treatment with Y-27632 is convenient because it acts rapidly and is easily washed out by replacing the media in the sample chamber with fresh media. Upon removal of Y-27632, the actin cytoskeleton begins to reform stress fibres and again takes on an aligned appearance (Fig 6B). As evidenced in the plot of the order parameter, we can see a loss of alignment immediately after addition of Y-27632 and an immediate recovery following washout (Fig 6C).

**Figure 6:**
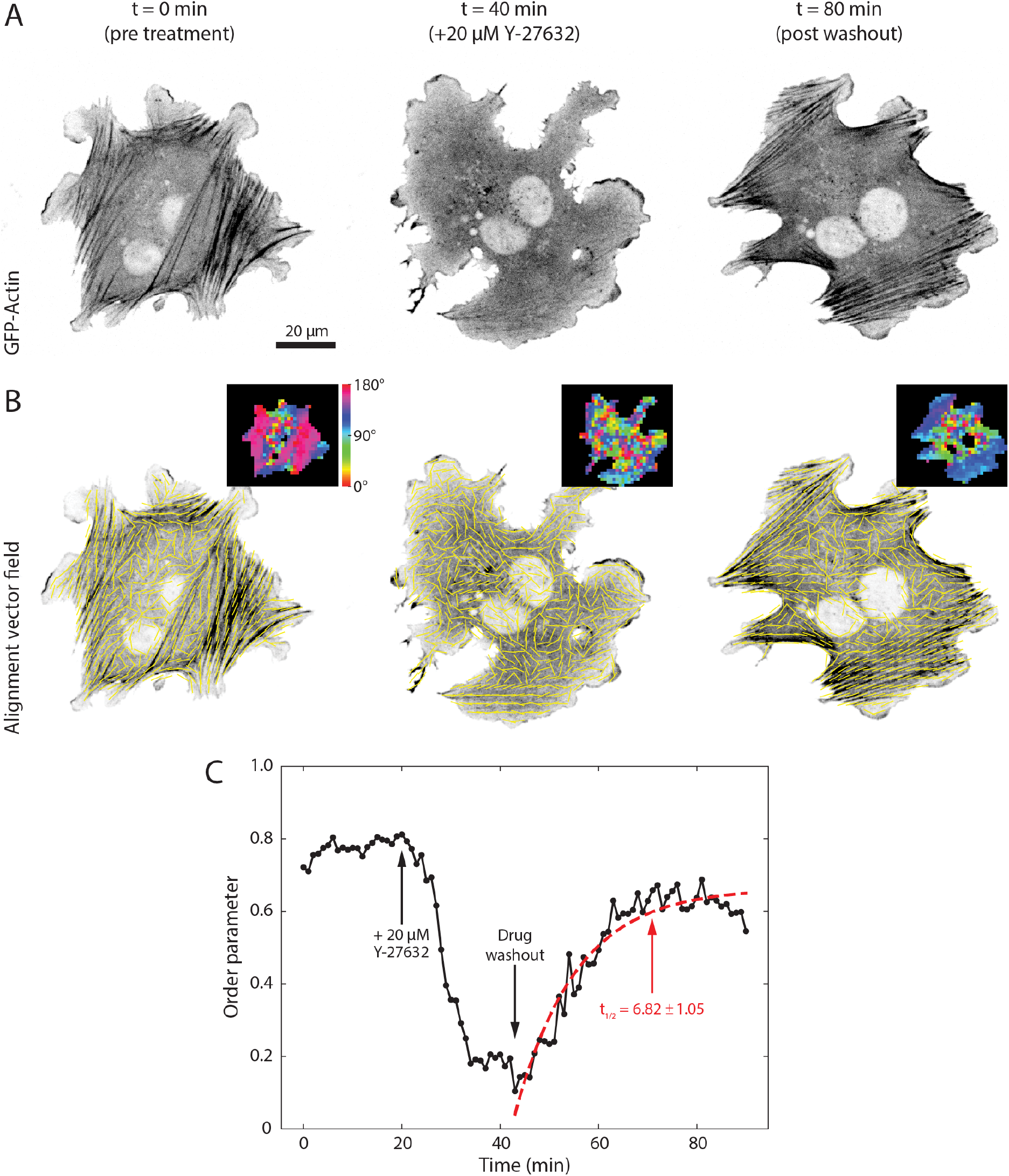
Measuring kinetics of alignment. **(A)** An NIH 3T3 fibroblast transfected with GFP-actin is shown before, during, and after washout of 20 *μm* Y-27632. **(B)** Local alignment was calculated for each frame in the time series using a window size of 17 *px* and an overlap of 50%. Yellow vectors indicate the local direction of alignment in the image while insets show the angle of orientation for each window. **(C)** Plot of the calculated order parameter over time for a neighbourhood radius of 2*x* vectors. Arrows indicate the time points of addition of Y-27632 and wash out of the drug. Red dashed line indicates the fit to Eq 7.

To measure the kinetics of this interaction we can fit the data to any relevant model. For the washout of Y-27632 we fit the post-washout data to a simple exponential recovery curve of the form

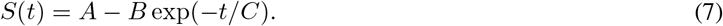

We can then define a *t*_1/2_ value, or the time it takes to reach half its maximum recovery value, as

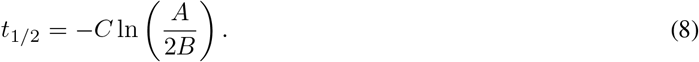

This results in a value of *t*_1/2_ ≈ 7 min for the Y-27632 washout experiment, consistent with previous reports (Aratyn-Schaus et al., 2011).

### 3.6 Comparing with other available algorithms

The performance of the algorithm presented in this work was tested against two other open-source tools widely used by the biology community, OrientationJ (Rezakhaniha et al., 2012) and TWOMBLI (Wershof et al., 2021). OrientationJ is a Fiji plugin based on image representation by structure tensors using real space, and evaluates orientation by the coherency metric, that indicates the degree of feature orientation with values ranging from 0 (random) to 1 (perfectly aligned) (Rezakhaniha et al., 2012; Clemons et al., 2018). Using this approach, it is possible to output either an alignment vector field (OrientationJ Vector Field) or a global coherency score for the image (OrientationJ Dominant Direction). TWOMBLI is a Fiji macro envisioned as a hybrid approach performing fibre segmentation followed by alignment analysis. Fibre segmentation is carried out with the Ridge Detection plugin (Lindeberg, 1998), while the alignment of the binarised image is subsequently measured via OrientationJ. In addition to the coherency metric as a measure of alignment and due to the segmentation step, TWOMBLI also allows the user to obtain further metrics that characterise the fibrillar features, such as number of branch and end points, density, curvature, and fractal dimensions.

We first compared the alignment vector field calculated with the three methods (FFT, structure tensor, and hybrid approach) for cell-derived matrices immunostained for fibronectin displaying different degrees of isotropy (Fig 7). When comparing the alignment vector field for images with strongly anisotropic fibrillar features, similar output was observed for the three methods (Fig 7A). When working with less aligned input, however, the FFT was able to better resolve the feature alignment in the isotropic regions compared to the other two algorithms (Fig 7B).

**Figure 7:**
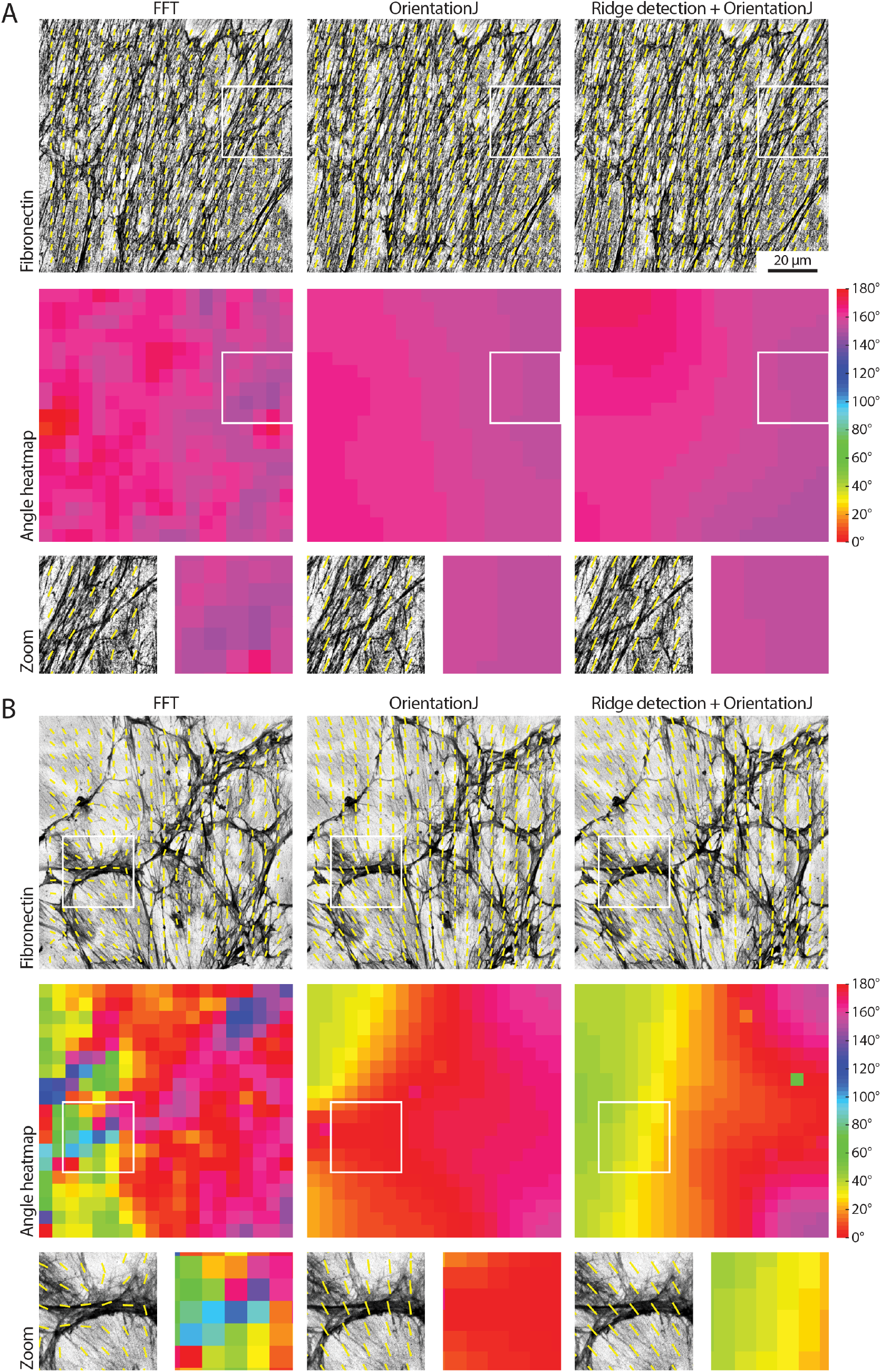
Comparing approaches to measure organisation in biological images. **(A)** A fibronectin image showing fibrillar features with strong alignment is analysed with the FFT approach presented in this work, with the Fiji OrientationJ Vector Field plugin, and with a combined approach using the Fiji Ridge Detection plugin followed by OrientationJ (mimicking the TWOMBLI hybrid approach). A window size of 100 *px* with 50% overlap was used, and the order parameter was calculated over neighbourhoods of 5*x* vectors (neighbourhood radius of 2*x* vectors). Both the alignment vector field and the angle heatmap are shown for each case; zooms for the areas highlighted by the white boxes are displayed. The three algorithms behave similarly. Scale bar, 20 *μm*. **(B)** The analysis of a more randomly organised fibronectin image is compared as in (A). It can be noted how the FFT returns more accurate alignment vectors for the more isotropic regions of the image (insets).

Two alignment metrics were analysed, the order parameter *S* (Eq 6) and the coherency; the first metric is the one used in this work, while the coherency is the standard output of both OrientationJ and TWOMBLI. The comparison dataset contained two samples of images, with isotropic (Fig 7B) and anisotropic (Fig 7A) features. All the tested algorithms were able to recognise the alignment difference in the data, however, the order parameter scores calculated with the TWOMBLI approach (fibre segmentation via Ridge Detection followed by alignment with OrientationJ) did not reach statistical significance (Fig 8A). Moreover, the order parameter values calculated with both OrientationJ and the hybrid approach were very close to the upper limit of the range (values equal to 1 signify perfect alignment). This suggests that these methods might not be reliable in detecting subtle differences between samples, due to their limited ability to resolve local isotropy (Fig 7). Global coherency obtained with the three methods was also compared, showing that all of them could successfully distinguish the global alignment differences in the two samples (Fig 8B).

**Figure 8:**
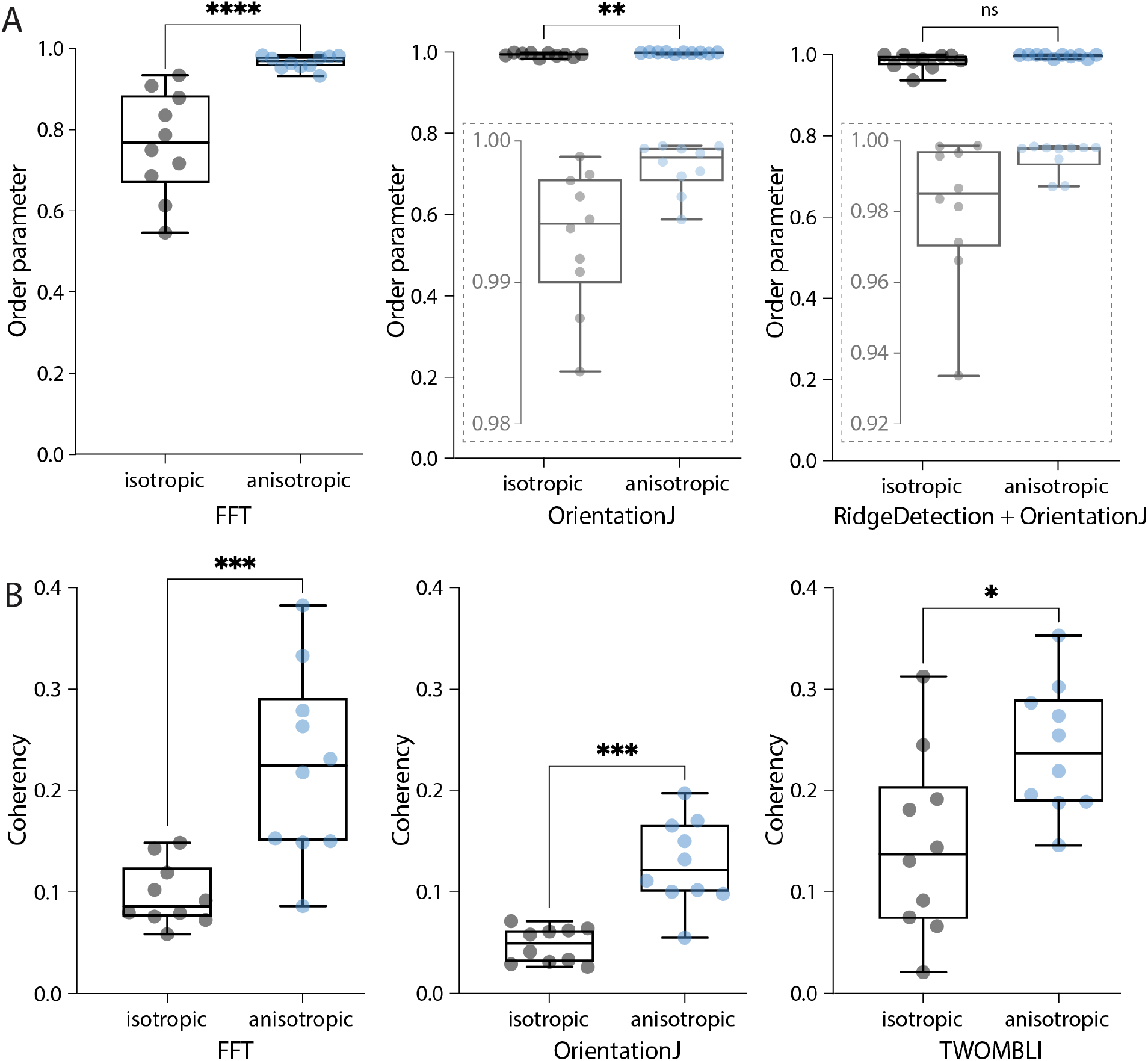
Comparing output metrics of different approaches to measure organisation in biological images. **(A)** Boxplots of the order parameters for the samples shown in Fig 7 calculated with the three methods, the FFT (as presented here), OrientationJ (Vector Field plugin), and the hybrid approach (Ridge Detection followed by OrientationJ Vector Field). Local windows of 100 *px* with 50% overlap and 5*x* vector neighbourhoods were considered. Insets represent zooms of the y-axis to better display distributions. Mann-Whitney two-tailed test (****p <0.0001, **p = 0.0039, ns = 0.063). **(B)** Boxplots of the global coherency calculated with the three tools, *AFT - Alignment by Fourier Transform* (as presented here), OrientationJ (Dominant Direction plugin), and TWOMBLI. Mann-Whitney two-tailed test (***p = 0.0002, ***p = 0.0002, *p = 0.0147).

Computational time is comparable between the FFT and OrientationJ (with a dedicated macro) with each image being evaluated in less than half a second, while it increases for TWOMBLI with about 7 s per image (due to the fibre segmentation step). While this difference might appear insignificant for small samples as the ones used in this context, it might become important should batch processing be required for larger sample sizes or more high-throughput applications. The FFT approach presented here can be easily used to address length scale analysis. While OrientationJ also offers the option of a length scale analysis (Vector Field plugin), this requires *ad hoc* macros and further post-processing to obtain a unique alignment score for each image for a given window size. TWOMBLI allows the user to obtain a wider range of metrics characterising the fibrillar features, thanks to its fibre segmentation step, but this comes at the cost of setting more parameters to start the analysis. The FFT performs best at resolving local alignment differences (Fig 7), and the available filters on window intensity and eccentricity make it straightforward to automatically exclude regions of the images. This avoids having to manually discard entire images containing unsuitable regions, as would happen with the other two methods. The pros and cons of each methodology should be taken into account depending on the application (Table 1); overall, the *AFT* approach demonstrated efficient performance and broad flexibility to different types of input images.

**Table 1:** A comparison of the performance and characteristics of the three analysis methods.

|  | **AFT** | **OrientationJ** | **TWOMBLI** |
| --- | --- | --- | --- |
| Computational time | low (1.8-4.8 s*) | low (4.8 s**) | high (72.4 s) |
| Length scale analysis | + | (+) | - |
| Additional metrics | - | (+) | + |
| Automated image filtering | + | - | - |

Computational time was measured on a 3.6 GHz Quad-Core Intel Core i7 machine with 32 GB memory for a 10-image sample. ‘+’: yes, ‘(+)’: yes, but requires further post-processing, ‘\*’: depends on window size; ‘\*\*’: with a dedicated macro.

### 3.7 Implementing a streamlined software for measuring alignment

Finally, we have developed a workflow named *AFT - Alignment by Fourier Transform* to automate image analysis using the approach described above. This includes the pipeline described in Section 3.1, the parameter search in Section 3.2, the analysis of length scale decay in Section 3.3, the filtering options in Section 3.4, and the extraction of organisation kinetics in Section 3.5. Open-source implementation is made available in both MATLAB and Python at https://github.com/OakesLab/AFT-Alignment_by_Fourier_Transform.

Some considerations regarding the choice of parameters follows. The user is requested to set three mandatory parameters in order to run the alignment analysis: the window size, the overlap, and the neighbourhood size (Section 3.1, Fig 1D and 2B). Each of these parameters will affect the calculated order parameter by changing the length scale of the alignment analysis. While default parameters are suggested, it is important to test different values on individual images to gain some insight on the optimal length scale over which to carry out this analysis. In addition to the mandatory parameters, a number of optional filtering features are available as described in Section 3.4 (Fig 5).

#### 3.7.1 Mandatory parameters

- *Window size*. The size of the local regions in the image to analyse. This value depends on the image resolution, the size of the fibrillar features to be examined, and the length scale of interest (Fig 1D). In practice, within a window one should be able to observe enough information to discern by eye an orientation of features. There is no hard-set lowest limit for the window size, but it should be noted that smaller windows will lead to more spurious vectors as the calculated FFT will be nearly circular (i.e., not skewed, Fig 1A) for regions where fibrillar features are not recognised. A visual check of the output images is always recommended during the parameter optimisation phase.
- *Overlap*. The overlap between adjacent windows (Fig 1D). Increasing the overlap, in conjunction with choosing smaller window sizes, allows for increasing the resolution of the analysis, with more sampled areas (i.e., vectors) within the field of view. A starting value of 50% is recommended.
- *Neighbourhood radius*. The length scale over which to compare neighbouring measures of alignment. This should be set depending on the length scale of interest. It is defined as the number of nearest neighbour vectors to be compared to a central reference in order to obtain an alignment score (Fig 2). Neighbourhood size is calculated as 2 * *neighbourhood radius +* 1.

#### 3.7.2 Optional parameters

- *Output images*. This parameter enables the ability to save the analysed images, consisting of the original image overlaid with the alignment vector field and the angle heatmap (Fig 1D). The colour scale for the angle heatmap is wrapped, with similar colours for 0° and 180°, as fibres are not considered polar (Section 3.1, Fig 1D).
- *Filter blank spaces*. Windows that contain little or no signal can be excluded from the analysis. When enabled a threshold on the mean window intensity can be set (Fig 5AB). Windows below this mean intensity are not analysed. Threshold values can be estimated by opening a sample image in Fiji, selecting regions considered background with similar size to the window size, and measuring their mean intensity. Expected values range from 0 (black) to 255 (white) for an 8-bit image.
- *Filter isotropic regions*. It is possible to exclude windows from the analysis should the eccentricity of their FFT be lower than a threshold (Fig 5CD). Values for the FFT eccentricity range from 0 (circular, isotropic) to 1 (elongated, anisotropic). To estimate this threshold, the analysis can be run iteratively for increasing values until the desired output is achieved (i.e., until the regions considered isotropic do not contain alignment vectors in the output images).
- *Masking*. By default, the whole image is being analysed. Alternatively, the user can input a folder containing binary masks of selected areas to be analysed for each input image (Fig 5E). A similar output as the one shown in Fig 5E could have also been obtained by filtering out the blank spaces, as the regions of interest have brighter signal than the background. However, if regions devoid of signal would also be present inside the cell outline, they would be disregarded as well. Therefore, a binary masking approach might be preferable in this case.

## 4 Discussion

Quantifying alignment of fibrillar features in biological images has gained wide interest as a possible biomarker in disease aetiology and progression (Ouellette et al., 2021). While tools are available for this aim (Püspöki et al., 2016; Liu et al., 2017), many are difficult to implement and fairly rigid on the length scale over which anisotropy can be interrogated. The method presented in this work exploits the representation of the image in the frequency domain paired with a custom window-based approach, offering a computationally efficient algorithm that can be easily applied to investigate local to global alignment. Decay in alignment scores with increasing distance can be evaluated to reveal the length scale at which fibrillar features remain anisotropic.

An open-source suite, *AFT - Alignment by Fourier Transform*, is presented, and its flexibility to a wide range of biological images is showcased through the examples. The *AFT - Alignment by Fourier Transform* tool is demonstrated to be robust for different imaging modes (fixed, live-cell, SHG, fluorescence), fluorescent probes (F-actin, fibronectin), and image resolutions (from single cells to tissue tilescans). Further applications for which this method could be used that were not shown in the present work could entail the evaluation of alignment across the depth of a sample (measuring alignment for each slice of a Z-stack), or the investigation of cytoskeletal architectures in reconstituted protein systems.

The FFT approach performance was comparable to both OrientationJ and TWOMBLI, with increased accuracy in resolving local isotropic regions. Its computational efficiency makes it a good candidate for analysing high volumes of data. Moreover, the availability of filters and masking options allow for this methodology to be applied to a wide range of microscopy images.

Excitingly, more advanced 3-dimensional imaging modalities are becoming available, with techniques being developed to achieve isotropic resolution. While currently the FFT approach is designed to work with 2D data, the algorithm could be adapted to evaluate alignment of 3D fibrillar features, and its low computational cost could be exploited for these more computationally intensive data.

## 5 Conclusions

Here we presented a methodology to measure fibrillar feature alignment in biological images. We created an open-source tool that can be used on a wide range of biological images. Its performance was compared against other common workflows, and it was shown to be computationally efficient without compromising on accuracy. It is easy to implement with a small number of analysis parameters, and allows for interrogating the length scale of fibre anisotropy.

## Author Contributions

Conceptualisation and Methodology, PWO; Software, SM, DBDF, PWO; Data acquisition, LDT, FK; Formal analysis and Data curation, SM, PWO; Writing SM, DBDF, LDT, FK, TS, BMS, PWO; Funding Acquisition and Supervision, TS, BMS, PWO.

## Software Availability Statement

The software implementations can be found at https://github.com/OakesLab/AFT-Alignment_by_Fourier_Transform, together with further documentation on how to run the code.

